# Noninvasive method for monitoring breathing patterns in monkeys

**DOI:** 10.1101/2022.08.31.505131

**Authors:** Jun Kunimatsu, Yusuke Akiyama, Osamu Toyoshima, Masayuki Matsumoto

## Abstract

Respiration is strongly linked to internal states such as arousal, emotion, and even cognitive processes and provides objective biological information to estimate these states in humans and animals. However, the measurement of respiration has not been established in macaque monkeys that have been widely used as model animals for understanding various higher brain functions. In the present study, we developed a method to monitor the respiration of behaving monkeys. We first measured the temperature of their nasal breathing, which changes between inspiration and expiration phases, in an anesthetized condition and estimated the respiration pattern. We compared the estimated pattern with that obtained by a conventional chest band method that has been used in humans and applies to anesthetized, but not behaving, monkeys. These respiration patterns matched well, suggesting that the measurement of nasal air temperature can be used to monitor the respiration of monkeys. Furthermore, we confirmed that the respiration frequency in behaving monkeys monitored by the measurement of nasal air temperature was not affected by the orofacial movement of licking to obtain the liquid reward. We next examined the frequency of respiration when they listened to music or white noise. The respiratory frequency was higher when the monkeys listened to music than the noise. This result is consistent with a phenomenon in humans and indicates the accuracy of our monitoring method. These data suggest that the measurement of nasal air temperature enables us to monitor the respiration of behaving monkeys and thereby estimate their internal states.

**Significance Statement:** While respiration is linked with internal processing, such as emotional and cognitive states, methods have not been established for physiological research on monkeys. We developed a novel method that obtained respiration signals by measuring the nasal air temperatures of behaving monkeys. Our method was able to continuously track the respiration pattern without distortions evoked by orofacial movements to lick the liquid reward. The respiratory frequency increased while listening to music than when listening to white noise for all monkeys. These results demonstrate that nasal air temperature measurements can be used to monitor the respiration patterns of aroused monkeys, allowing us to understand their internal state. This will be useful for investigating the underlying neuronal mechanism of neuropsychiatric disorders using monkeys.

## Introduction

While humans can verbally explain their internal states such as emotions and feelings, animals cannot. However, it is possible to estimate the internal states of animals by monitoring their behavior and physiological parameters. Since an altered internal state is a remarkable characteristic of psychiatric disorders such as depression and schizophrenia (Boiten et al., 1994), techniques to monitor physiological parameters have made significant contributions to animal model studies on mechanisms underlying these disorders (Lang et al., 2000; Rudebeck et al., 2014). In previous studies using animals, physiological parameters, including heart rate (Maeda et al., 2018; Wascher, 2021), pupil diameter (Suzuki et al., 2016; Maeda et al., 2018; Kuraoka and Nakamura, 2022), and skin temperature (Nakayama et al., 2005; Kuraoka and Nakamura, 2022), were measured to estimate emotion and arousal. Among them, respiration is a well-measured physiological parameter, the pattern of which changes depending on internal states such as anxiety and stress (Suess et al., 1980; Boiten, 1998; Homma and Masaoka, 2008). In contrast to other physiological parameters that are involuntarily regulated, respiration is controlled not only involuntarily but also voluntarily. Thus, this physiological parameter is thought to reflect richer internal information than the other parameters. Consistent with this idea, brain regions that affect respiration are widely distributed over the cortical and subcortical areas. Several regions in the medulla are known to directly control respiration (Del Negro et al., 2018). In addition, respiration is modulated by neural oscillations in the limbic brain regions, such as the amygdala and hippocampus, that are involved in emotion (Frysztak and Neafsey, 1991; Zelano et al., 2016). Human studies have also reported that neural oscillations in the prefrontal cortex, a center of cognitive processes, are synchronized with the rhythm of voluntary respiration (McKay et al., 2003). These data suggest that brain activities regulating not only emotion but also cognitive processes affect respiratory control, and thereby respiration measure is likely to be useful to estimate these internal states.

Behavioral and neuroscience studies on emotion and cognitive processes have used macaque monkeys as experimental animals due to their similarities to humans in behavioral capacities and brain architecture. However, these studies did not monitor their respiration because there is no established method to monitor respiration in behaving monkeys. Body and air movements evoked by respiration (e.g., chest movement) were measured to monitor respiration in humans (Porges and Byrne, 1992) and rodents (Grimaud and Murthy, 2018). The measuring equipment (e.g., chest band) was directly attached to the body. These conventional methods are hard to apply to behaving macaque monkeys because they have dexterous fingers like humans and often remove the measuring equipment. To monitor the respiration of behaving macaque monkeys, the present study developed a method to measure the temperature of their nasal breath (nasal air) that changes between inspiration and expiration phases. The equipment did not touch any part of the body so that the monkeys were not disturbed, and was attached to a location where the monkeys could not approach it. We successfully reconstructed the respiration pattern of behaving monkeys from the measured nasal air temperature. This methodology enables us to estimate internal states of behaving macaque monkeys and facilitates research on neuronal mechanisms underlying respiration, motivation, and cognitive processes in primates.

## Materials and Methods

### Animal preparation

Two female Japanese monkeys (monkeys P and M, 9 and 5 years old, 7.0 and 6.2 kg, respectively) and a male rhesus monkey (monkey S, 9 years old, 5.9 kg) were used in the experiment. All procedures for animal care and experimentation were approved by the University of Tsukuba Animal Experiment Committee (permission number: 19-006) and were carried out according to the guidelines described in the Guide for the Care and Use of Laboratory Animals published by the Institute for Laboratory Animal Research. Initially, the monkeys were habituated to sitting on a primate chair. Using sterile procedures and general isoflurane anesthesia, a head holder was implanted, which was embedded in dental acrylic covering the top of the skull and firmly anchored using plastic screws. Analgesics were administered during each surgery and for several days afterward. The experimental sessions began after the monkeys had fully recovered from the surgery.

### Respiration and licking behavior monitoring

To monitor the respiration of the monkeys, we measured the temperature of nasal air, which changes between inspiration and expiration phases, using a thermosensor (Figure 1A; Model BAT-12R, Physitemp) not only in the anesthetized condition (medetomidine and midazolam) but also in the awake condition. Nasal air temperature is maintained uniform by the ambient temperature during the inspiration phase but increases during the expiration phase in which warm air is expired. We also measured changes in the monkey’s chest diameter using a movement sensor (TN1132/ST, AD Instruments) in the anesthetized condition. During the experiment, the monkey sat in a primate chair in a sound-attenuated room, and its head was fixed using the head holder. We measured the nasal air temperature with the thermosensor that was located at three different positions: 5 mm outside, 0 mm, and 5 mm inside from the entrance of the nasal cavity. When we measured the temperature in the awake condition, the thermosensor was located inside. Licking movements were measured with a vibration sensor (AE-9922, NF Corporation) that was attached to the spout for reward delivery. These analog data were collected using a data acquisition system (Omniplex, Plexon).

**Figure 1.**
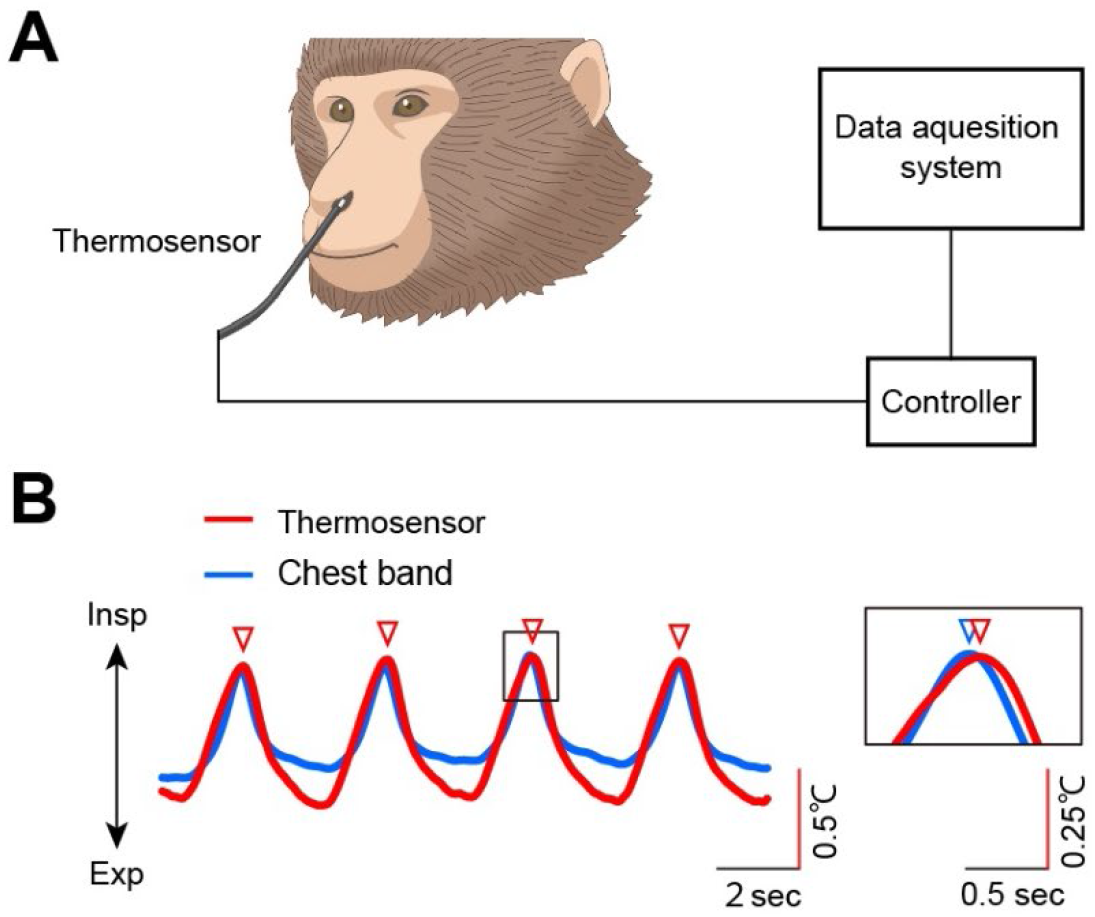
Procedure to monitor the respiration of the monkey using a thermosensor. (A) Thermosensor is placed in the nasal cavity and measures air temperature change with respiration. (B) Left, a representative trace of the respiratory signal from monkey P was monitored with a temperature sensor (red) and movement sensor (blue). The nasal air temperature increased and decreased with the expiration and inspiration, respectively. The red triangles indicate the peaks of the respiratory phase waveform monitored by the temperature sensor. Right, enlarged view of the area indicated by the solid box. The blue triangle indicates the peak of the respiration signal monitored with the chest band method.

### Sound stimulation and reward delivery

In the awake condition, sound stimuli and a liquid reward were presented to the monkeys. These sound stimulation and reward delivery were controlled with a personal computer running the MonkeyLogic Matlab toolbox (Asaad and Eskandar, 2008). We used three different tempos of music for sound stimulation (“Master of Puppets” by Metallica, 212 beats/min; “Berceuse” by Chopin, 85 beats/min; and “Can’t stop the feeling” by Justin Timberlake, 113 beats/min; 65.0-70.0 dB SPL). The introductions of the music were removed to match their volume levels. We also used three pieces of white noise (40.0 dB SPL) to measure the resting respiration frequency. Each music and white noise were presented for 3 min with a 1-min interval. Figure 3A shows the sound stimulation schedule of one recording session. The recording session was executed once a day and was repeated for three days. The music and white noise were presented using a speaker that was located 50 cm away from the monkey’s face. A liquid reward (0.2 mL) was delivered randomly (once in 30 sec) during the sound stimulation.

### Analysis

The respiration and licking data were analyzed offline using a commercial software (Matlab 2018b, Mathworks). We calculated the chest diameter by integrating the chest movement signals. To calculate the frequency of respiration, we identified peaks of the respiratory waveform that corresponded to the time when the phase of respiration changed from inspiration to expiration (Figure 1B). To count the number of licks, we detected the onset of each lick as the time when the licking signal measured by the vibration sensor reached a threshold that was manually adjusted for each day (Figure 3B).

## Results

### Accuracy of the respiration signal monitored by nasal air temperature

To develop a noninvasive respiration monitoring system for macaque monkeys, the present study focused on the temperature of nasal air that changes between inspiration and expiration phases and measured it with a thermosensor located at the entrance of the nasal cavity (Figure 1A). This thermosensor detected the resistance of the material at its tip that changed with the temperature, with a high temporal resolution (100 ms). We first applied this system to the three anesthetized monkeys (monkeys P, M, and S) and observed an oscillation of nasal air temperature that could coincide with the alternation between inspiration and expiration phases (red curve in Figure 1B). The peaks of the oscillating temperature signal (red triangles in Figure 1B) were corresponding to the times when the phase of respiration changed from inspiration to expiration. To examine how accurately the temperature signal reflected the pattern of respiration, we simultaneously monitored the respiration of the anesthetized monkeys by a conventional chest-band method that had also been used in humans and applicable to anesthetized monkeys (blue curve in Figure 1B). We observed that the temperature and respiration signals were well synchronized. However, the temperature signal obtained by the thermosensor was slightly delayed compared to the respiration signal obtained by the chest band (see the enlarged panel in Figure 1B).

The position of the thermosensor in the nasal cavity may have caused this delay. To find the best position, we measured the nasal air temperature at three different positions—5 mm inside, 0 mm, and 5 mm outside the entrance of the nasal cavity. We then calculated the delay in the temperature signal compared to the respiration signal calculated by the chest band for each position (i.e., the time difference between the peaks of the temperature signal and those of the respiration signal) (Figure 2A). We observed that the delay in the temperature signal decreased when the thermosensor was located inside the nostril. Although a significant decrease in the delay was observed only in one of the three monkeys (regression analysis; monkey M, r = -0.49, p = 0.01; monkey P, r = -0.13, p = 0.51; monkey S, r = -0.28, p = 0.11), this trend was significant as a whole (regression analysis; all monkeys, r = -0.23, p = 0.03). Hence, the thermosensor should be located on the inside rather than the outside or entrance of the nostril to reduce the temperature signal delay.

**Figure 2.**
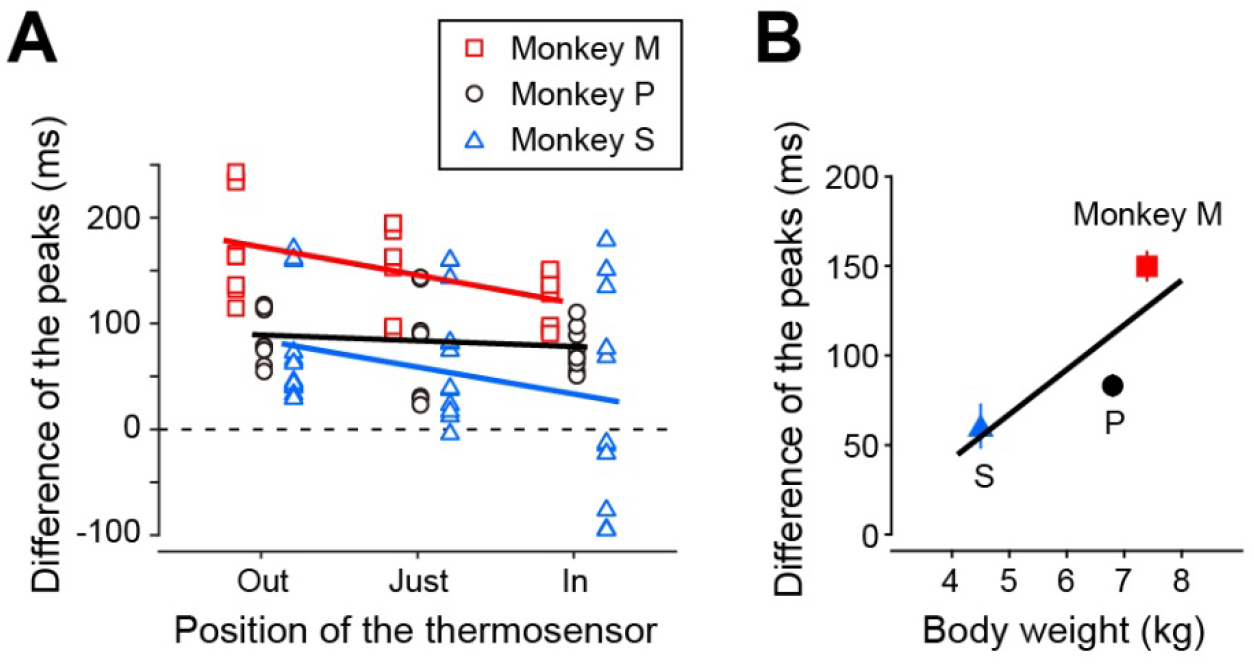
Differences in the respiration signals from the two measurements. (A) Each plot indicates the difference in the peak timing of respiration signal monitored by the thermosensor and chest band in three monkeys (red square, monkey M; black circle, monkey P; blue triangle, monkey S). The temperature was measured from three positions [-5 mm (out), 0 mm (just), and 5 mm (in) from the entrance of the nasal cavity] in the three monkeys. Each line indicates the regression line. (B) The differences in peak timing depend on the body weight. The mean of the peak timing recorded from the three probe positions was plotted.

Although the delay of the temperature signal obtained by the thermosensor compared to the respiration signal calculated by the chest band was small (0 to 100 ms in the “inside” condition), researchers that need a fine temporal resolution may have to correct this delay. A factor that could influence this delay is the body weight of the monkeys because the volume of airflow evoked by respiration changes depending on the weight. Figure 2B shows the relationship between the temperature signal delay and body weight. We observed that the delay tended to increase as the body weight increased (regression analysis; r = 0.47, p = 2.58 × 10^−6^). Thus, correction values and correction coefficients to correct the delay of the temperature signal obtained by the thermosensor need to be determined in accordance with the body weight.

### Respiration monitored by nasal air temperature in behaving monkeys

The results indicate that the respiration of anesthetized monkeys can be monitored by measuring the nasal air temperature. We next applied this method to behaving monkeys. The nasal air temperature of the three monkeys was measured while sound stimuli and a liquid reward were presented. Three music and white noise were sequentially presented for 3 min each with a 1-min interval (see Figure 3A for the schedule).

Similar to the anesthetized condition, we observed an oscillation of nasal air temperature that reflected the alternation between inspiration and expiration phases during the presentation of the music (lower curve in Figure 3B). We examined whether the temperature signal was affected by orofacial movements to lick the liquid reward. Liquid rewards are commonly used in neuroscience and behavioral studies on macaque monkeys. Therefore, to disseminate our method in these fields, it is useful to show that respiration patterns monitored by our method are not distorted by orofacial movements to lick the liquid. The upper curve in Figure 3B shows a typical example of the licking signal measured by a vibration sensor that was attached to the spout for reward delivery. To quantify the effect of licking on the temperature signal, we aligned the signal at the onset of lick events (shown by inverted triangles in Figure 3B). We observed that the temperature signal did not change before and after the lick events (Figure 3c). This result suggests that respiration patterns monitored by nasal air temperature are not distorted by orofacial movements to lick the liquid and that our method can be used in conjunction with liquid rewards.

**Figure 3.**
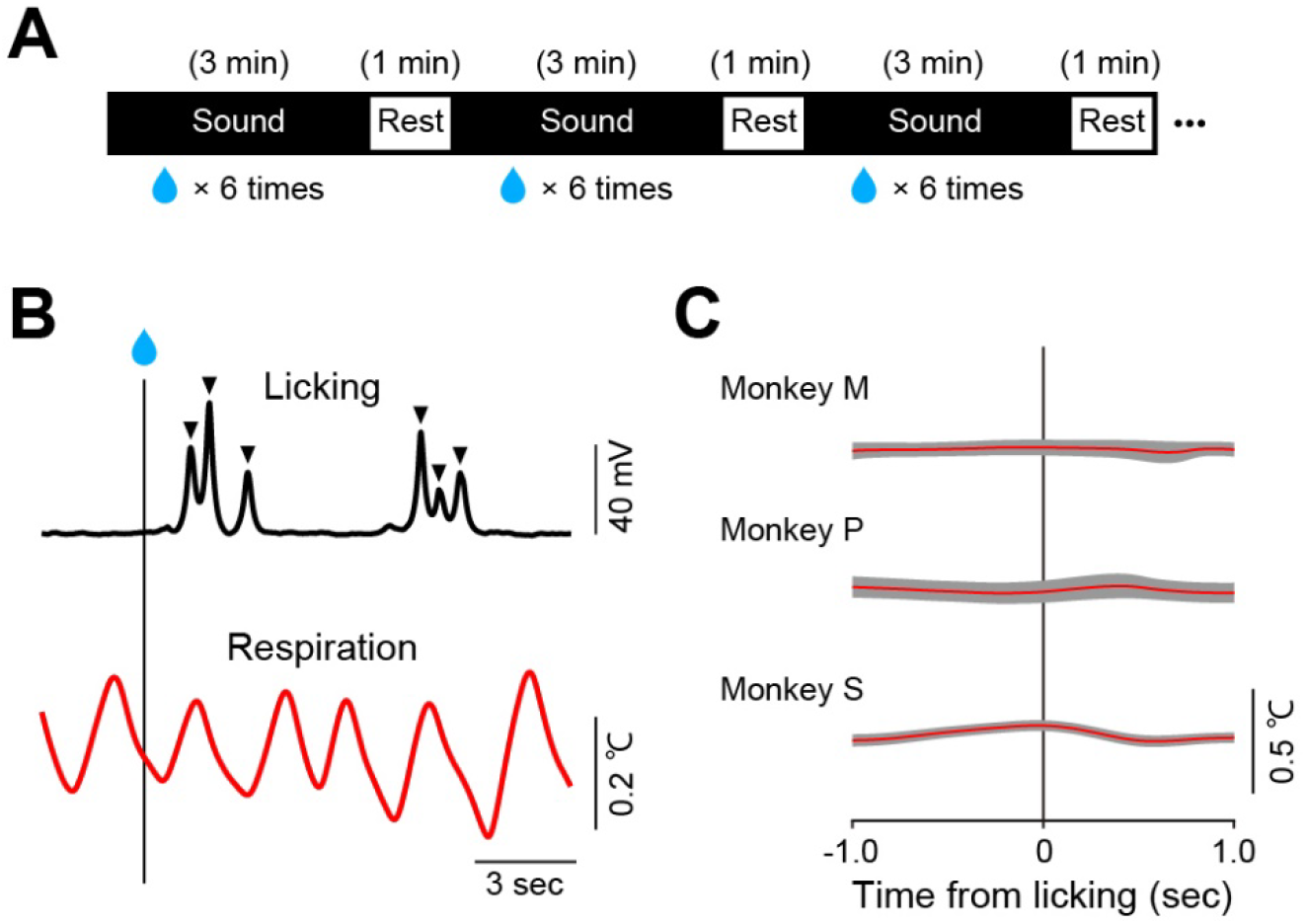
Effect of the licking on respiration monitoring. (A) Schedule of the music stimuli and reward delivery. One sound stimulus was chosen pseudo-randomly from three candidates and was presented for 3 min at 1 min intervals. The monkeys received six rewards irregularly during the sound stimulation. (B) The example of the licking (above) and respiration (bottom) after rewarding monkey M. Triangles indicate the timing of single licking. (C) Mean and SD of the temperature aligned on the licking during one session (monkey M, 84licks; monkey P, 214; monkey S, 90).

### Rhythmic auditory stimuli changing the frequency of respiration

Previous studies have reported that, in humans, while rhythmic auditory stimuli such as music are presented, the frequency of respiration changes depending on the rhythm (Bernardi et al., 2006; Salimpoor et al., 2009). Here we tested whether the same phenomenon occurs in monkeys. We monitored the respiration of the three monkeys by measuring their nasal air temperature while they listened to the three types of music with different tempos (fast, 212 beats/min; medium, 113 beats/min; and slow, 85 beats/min) as well as white noise, and compared the respiration frequency during the presentation of the rhythmic auditory stimuli (i.e., music) with that of the nonrhythmic auditory stimulus (i.e., white noise). Figures 4A and 4B show the inter-peak interval of the measured temperature signal, which reflects respiration frequency, of monkey S in the one-day experiment. The inter-peak interval during music presentation (including three music type data) was significantly smaller than that during white noise presentation (Figure 4A; unpaired t-test, *p* = 2.02 × 10^−5^). However, the inter-peak interval was not significantly different among the three types of music (Figure 4B; ANOVA, *p* = 0.32).

**Figure 4.**
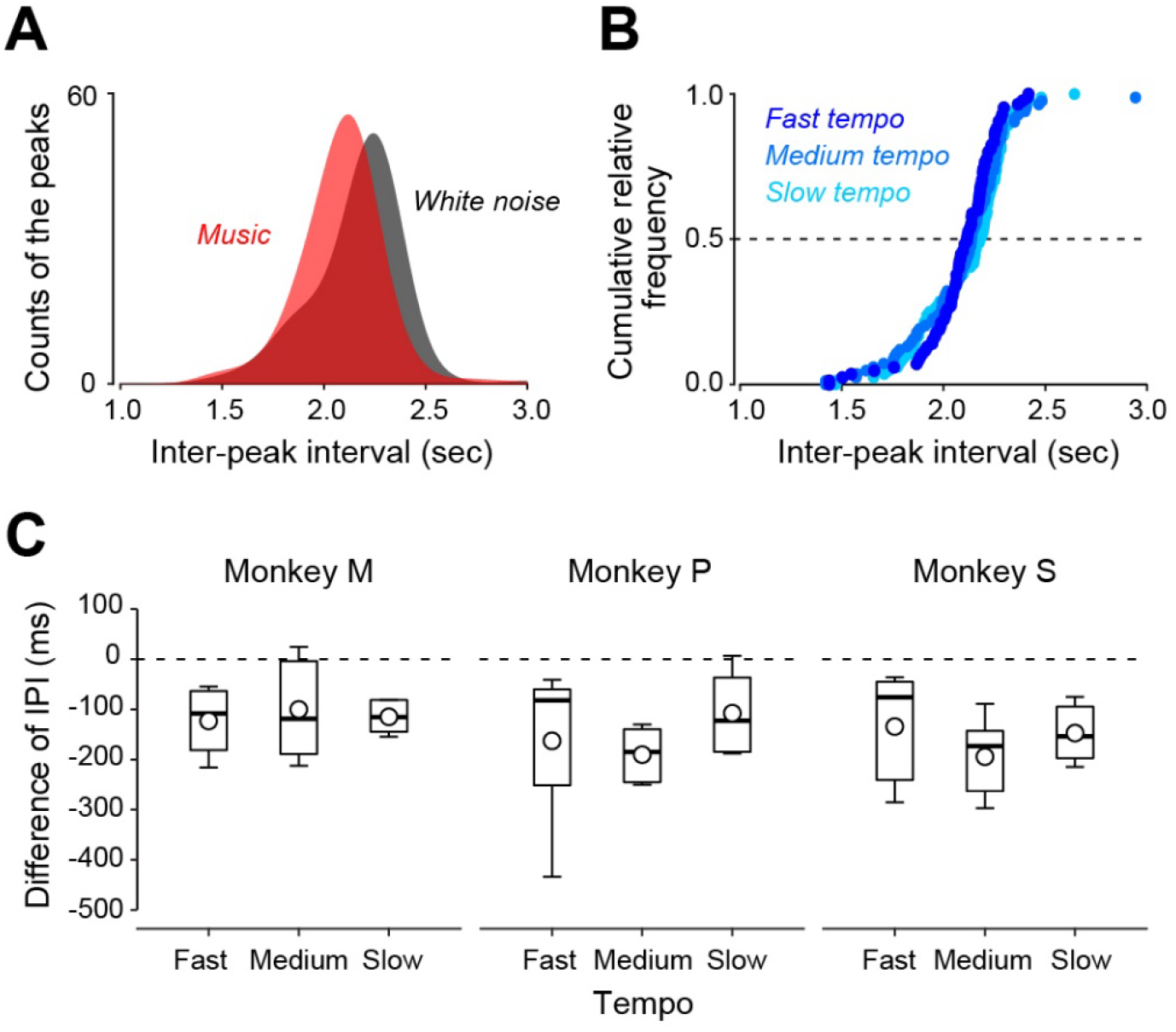
Change in respiration caused by rhythmic auditory stimulus. (A) The inter-peak intervals (IPI) when monkey S received the rhythmic auditory stimulus (red) and auditory white noise stimulus (grey) in one session. (B) The IPI while the monkey received the different tempos of auditory stimulus (fast, 212 beats/min; medium, 113 beats/min; slow, 85 beats/min). The data were the same as (A). (C) The differences in the IPI during fast, medium, and slow tempo of the music and white noise presentation in each monkey. The box-whisker plots indicate the median, quartiles, and range of the difference in IPI. Circles indicate the mean of the data.

We repeated the same experiment five times for each monkey and compared the inter-peak interval between each music and white noise (Figure 4C). We observed that many comparisons exhibited a significant difference (Figure 4C; paired t-test; monkey P, fast music *vs* noise, *p* = 0.09, medium music *vs* noise, *p* = 1.6 × 10^−3^, slow music *vs* noise, *p* = 0.04; monkey M, fast music *vs* noise, *p* = 0.02, medium music *vs* noise, *p* = 0.10, slow music *vs* noise, *p* = 1.6 × 10^−3^; monkey S, fast music *vs* noise, *p* = 0.06, medium music *vs* noise, *p* = 5.9 × 10^−3^, slow music *vs* noise, *p* = 5.0 × 10^−3^). Yet, the inter-peak interval was not significantly different among the three music in any of the monkeys (ANOVA, monkey M, *p* = 0.98; monkey P, *p* = 0.60; monkey S, *p* = 0.93). Taken together, these results suggest that the respiration of macaque monkeys is accelerated by rhythmic auditory stimuli but is not influenced by the tempo.

## Discussion

In this study, we succeeded in monitoring the respiration pattern of behaving macaque monkeys, which reflects their internal states such as emotion and cognition, by measuring the temperature of their nasal airflow. This noninvasive method is simple but has several features that make it useful in various physiological experiments. First, measuring the nasal air temperature allowed us to obtain accurate respiration patterns that were only slightly delayed from those obtained by the conventional method using a chest band (Figure 1). Second, our method was able to continuously track the respiration pattern without distortions evoked by orofacial movements to lick the liquid reward (Figure 3). Third, our method does not disturb monkey behavior because the thermosensor was non-contact. We succeeded to recorded the monkey’s natural physical response to the change in the respiration frequency due to rhythmic auditory stimulus (Figure 4). Thus, our respiration monitoring method would be a strong tool for estimating the internal state of behaving macaque monkeys.

Other methods have been used to monitor the respiration pattern of humans and animals (Grimaud and Murthy, 2018). Respiration is led by volume changes in the lungs induced by respiratory muscle activity. Therefore, the electromyography of inspiratory and expiratory muscles can be used to monitor respiration patterns (Murphy et al., 2001). Pleural pressure, which increases during the inspiration phase and decreases during the expiration phase, is also a physiological parameter that reflects the respiration pattern, and therefore can be used as its monitor. However, the measurements of respiratory muscle activity and pleural pressure require sophisticated surgical skills to implant electrodes and pressure sensors, respectively. It is difficult to apply these invasive measurements to behaving macaque monkeys because they can detach these probes with their dexterous fingers. In humans, measuring the chest diameter, the carbon dioxide of respiratory airflow, and the breathing flow rate are standard noninvasive methods to monitor respiration patterns (Porges and Byrne, 1992; Wientjes, 1992; Al-Khalidi et al., 2011). These methods provide accurate respiration signals, but the obtained signals may be distorted and may not reflect natural respiration, especially in behaving animals, because the instruments necessary for these methods, such as chest band and plastic facial mask, are directly attached to the body and disturb their behavior.

This study compared the respiratory signals from the chest diameter and nasal air temperature of anesthetized monkeys and revealed that both respiration signals correlated. We also found that the time difference between these signals is related to body weight (Figure 2). This might be because the volume of airflow evoked by respiration changes depending on weight. Our study also had other limitations. First, it is difficult to obtain quick changes in respiration signals through our method because temperature changes are gradual. Furthermore, there is a time lag (approximately 100 ms) in the inhalation onset detection between the methods presented here (Figure 2). The main reasons for the delay may be the time constant of the thermosensor and the nasal air temperature affecting signal detection. The time lag is affected by the position of the sensor in the nasal cavity (Figure 2A). Thus, although measuring the nasal air temperature provides adequate information to monitor the respiration rate and tidal volume, it does not reflect the quick changes in respiration. To examine the rapid response of the internal state, it might be useful to measure the pupil diameter (Suzuki et al., 2016; Kuraoka and Nakamura, 2022) or skin temperature (Nakayama et al., 2005; Kuraoka and Nakamura, 2022) concurrently with respiration. Second, our method uses the head-fix condition, which is commonly used in primate neurophysiology experiments, to enable stable measurements. Although we can obtain respiration signals in a head-free condition if the sensor is attached to the nasal area (ex. by using surgical tape), when the monkeys move their heads, there may be some signal changes. Hence, measuring the temperature of nasal airflow is suitable for respiratory monitoring in head-fixed aroused monkeys.

Respiration has a strong link with the internal state, such as cognitive and emotional processing. In humans, the respiration frequency changes depending on the tempo of the music regardless of the habituation, i.e., it increases with fast tempo and decreases with slow tempo (Haas et al., 1986; Bernardi et al., 2006; Gomez and Danuser, 2007; Salimpoor et al., 2009) but see (Sato et al., 2012). This suggests that faster rhythms may concentrate the attention and pauses or slower rhythms may induce relaxation. Conversely, our study revealed that the respiration rate in monkeys increased with both slow and fast music tempos (Figure 4C). The arousal in the animals may have occurred because of the musical sound, irrespective of the tempo. Alterations in respiration patterns are also found in psychiatric disorders such as anxiety and schizophrenia, neurological disorders, and aging (Boiten et al., 1994; Caldirola et al., 2004; Ramirez et al., 2013). For example, irregular respiration rates were found in patients with anxiety disorders, particularly when they were angry (Stevenson and Ripley, 1952). Research on the mechanisms of these emotion- and cognitive-respiration relationships is still limited. Tracking the connection between respiration patterns and performance in cognitive tasks and neurophysiological correlates provides an invaluable objective measure of the internal state. Subsequently, our method is a strong and useful tool for investigating the underlying neuronal mechanism of psychiatric disorders using non-human primates.

## Acknowledgments

We thank H. Yamada and T. Koganezawa for the valuable comments on the study; M. Isoda, A. Uematsu, T. Ninomiya, and A. Noritake for the technical advice; S. Nishino and Y. Narita for the animal care; and Y. Suwa and Y. Yabana for their administrative help. The Japanese monkeys were provided by the National Bio-Resource Project.

